# FADS and semi-rational design modified T7 RNA polymerase reduced dsRNA production, with lower terminal transferase and RDRP activities

**DOI:** 10.1101/2024.05.23.595468

**Authors:** Qiongwei Tang, Sisi Zhu, Nannan Hu, Sainan Yin, Yuhong Yang, Yigang Teng, Dongliang Song, Xiang Liu

## Abstract

T7 RNA polymerase (T7 RNAP) is the preferred tool for *in vitro* transcription (IVT), it synthesizes mRNA while accompanied by the generation of dsRNA by-products. This undesirable dsRNA triggers immune stress responses, compromises therapeutic efficacy, and raises safety concerns. To evolve T7 RNAP for reduced dsRNA, we pursued two complementary strategies. Firstly, the FADS (fluorescence-activated droplet sorting) based on molecular beacons was used to screen random libraries with diversity exceeding 10^5^. Secondly, we constructed several single-site saturated libraries to facilitate the transition of T7 RNAP from the initiation to the elongation conformation. These libraries were screened using the traditional microplate-based dual-probe screening technique. Both approaches identified two dominant variants: Mut1 (V214A) and Mut7 (F162S/A247T) from FADS, Mut11 (K180E) and Mut14 (A70Q) from saturated libraries. Furthermore, the combinatorial mutant Mut17 (A70Q/F162S/K180E), generated via DNA shuffling, exhibited significantly reduced dsRNA production compared to the wild-type under various conditions, ranging from 0.18% to 1.80%, with a minimum value of 0.5 pg/μg. Cell experiments confirmed that variants generated capped-mRNA with similar quality and quantity to the wild-type, while significantly reducing immune stress response in cells. These results indicate the compatibility and broad potential applications of these mutations. We then observed a close correlation between the production of dsRNA and the activities of T7 RNAP in terminal transferase and RDRP. Particularly, the terminal transferase activity appears to play a critical role in dsRNA generation. These findings align with the mechanism of dsRNA formation during IVT and provide new screening criteria for further evolution of T7 RNAP.

## Introduction

In recent years, mRNA-based drugs have garnered significant attention and attracted research focus. Particularly, mRNA vaccines, as a relatively new vaccine category, have demonstrated promise as an immunotherapy strategy[1,2,3]. Compared to traditional vaccines, mRNA offers several advantages. One crucial advantage is that the design and production of mRNA vaccines are independent of sequences. Different mRNA vaccines can utilize similar production and purification processes, enabling rapid and flexible development, and cost savings[4]. Additionally, mRNA vaccines can stimulate immune cells to secrete cytokines like tumor necrosis factor-α (TNF-α) and interferon-α (IFN-α), activating enduring adaptive immune responses without additional adjuvants[5]. To ensure clinical application, multiple steps are involved in mRNA production, including mRNA generation, DNA digestion, purification, and downstream processing. RNA polymerase facilitates *in vitro* transcription (IVT) to produce mRNA from linear dsDNA (double- stranded DNA) templates, the products typically containing the target protein’s open reading frame (ORF), UTR (untranslated region), 5’-cap, and 3’-poly(A) tail. Downstream purification processing involves precipitation and HPLC to remove impurities like redundant raw materials, DNA templates, proteins, and dsRNA (double-stranded RNA) by-products from transcription[6,7].

dsRNA, a pathogen-associated molecular pattern (PAMP), activates pattern recognition receptors in cellular compartments. Its recognition triggers the production of type I interferon, upregulating the expression and activity of protein kinase R (PKR; also known as EIF2AK2) and 2’-5’-oligoadenylate synthetase (OAS). Consequently, translation of intracellular proteins is suppressed, while degradation occurs for mRNA and ribosomal RNA[8,9,10]. To ensure the safety and efficacy of mRNA vaccines, it is necessary to remove dsRNA from the mRNA products. Currently, HPLC purification is an effective method for this purpose[8]. However, it increases time and costs, posing challenges for large-scale production and potentially leading to RNA degradation or reduced yield.

The bacteriophage T7 RNA polymerase (T7 RNAP) is the most commonly used enzyme for IVT. The 99 kDa protein consists of an N-terminal domain (residues 1-267) and a C-terminal domain (residues 268-883)[11–14]. T7 RNAP is known for its simplicity and ability to catalyze RNA synthesis with high yield and fidelity *in vitro*. However, during the transcription process, T7 RNAP can produce by-products such as abortive short RNAs, prematurely terminated RNAs, and 3’-extended RNAs, which creates favorable conditions for the formation of dsRNA[11,14–22].

The transcription of T7 RNAP involves three stages: initiation, elongation, and termination. During initiation, T7 RNAP recognizes specific promoters, opens the DNA double helix, and guides the template strand into the active site. The formation of this initiation complex marks the beginning of transcription. When the nascent mRNA reaches 2-6 nt, a conformational change occurs in the N-terminus of T7 RNAP. Multiple helices (D, E, F, G, I, J) in the promoter- binding region rotate, despite maintaining contact with the promoter. Simultaneously, helix C (residues 28-71) and subdomain H (residues 151-190) undergo a process of refolding. Helix C2 (residues 46-55) stacks onto the helix C1 (residues 28-41), resulting in the fusion of the original two short helices into a continuous long helix. Additionally, the loop (residues 173-186) within subdomain H transforms into an α-helix. These conformational rearrangements lead to the elimination of the promoter-binding site, expansion of the active pocket, and opening of the product RNA exit channel. Once the transcript reaches 10-12 nt, the conformational transition is complete, and the DNA promoter is released, marking the start of the elongation phase[11,13,14].

T7 RNAP continues to transcribe along the DNA template until it encounters a specific terminator or reaches the end of the linear DNA template, resulting in the generation of full-length RNA, known as run-off RNA[11,14–20]. However, premature termination during elongation can lead to incomplete RNAs, while non-template-dependent additions at the 3’-end of run-off RNA can generate 3’-extended products[21,22]. Additionally, the transition from initiation to elongation conformation is a challenging process that often leads to the accumulation of short single-stranded RNA, known as abortive RNA[11,14–17]. With the assistance of RDRP (RNA-dependent RNAP) activity of T7 RNAP, these 3’-extended products and abortive RNAs contribute to the formation of loopback dsRNA or complementary pairing-dependent dsRNA, leading to immunostimulatory effects in host cells[21–25].

Fluorescence-activated droplet sorting (FADS) is an emerging microfluidic-based screening technology that has gained popularity in the directed evolution of enzymes and strain selection. It offers several advantages, including ultra-high throughput, high sensitivity, and broad applicability[26,27]. FADS operates on a mechanism similar to fluorescence- activated cell sorting (FACS), where the device detects fluorescence of each droplet instead of cell and activates the electrodes to divert positive droplets into collection channels[28]. A typical droplet in FADS contains a lysing reagent, fluorescent substrate, and a microbial cell that overexpresses a mutant of the target enzyme. After sorting, the DNA within positive droplets can be amplified through PCR and subsequently used for sequence analysis to determine the variation types and abundance of dominant variants[29]. While in certain cases, secreted proteins or special substrates can directly pass through the cell membrane to activated fluorescence of droplet, without the need for cell lysis. This method enables direct culture of cells within positive droplets, facilitating the amplification of target clones with desired mutations[29].

Compared to traditional screening techniques, FADS provides a remarkable improvement in throughput, exceeding 10^3^-fold, and significantly reduces reagent consumption by more than 10^6^-fold. This is achieved due to the ultra-high throughput of FADS, which can reach 10⁶-10⁸ droplets per day while allowing for a reduction in reaction volumes within droplets to the picolitre range[30]. FADS has proven effective in directed evolution of various enzymes by directly modifying substrates with fluorescent groups or coupling with other reactions, including proteases, polymerases, RNase, glycosidases, and dehydrogenases[31]. The emergence of similar technologies, such as AADS (Absorbance-activated droplet sorting), BADS (BDD electrode-activated droplet sorting), and MADS (Mass-activated droplet sorting), has greatly expanded the application of high-throughput screening in enzyme evolution[32]. Moreover, there is great anticipation regarding the combination of these high-throughput screening technologies with IVTT (*In vitro* transcription and translation). IVTT is considered a valuable approach for expressing enzymes with cytotoxicity while also eliminating protein expression variations caused by the host[33].

We propose that preventing the production of dsRNA during IVT is a more efficient and economical approach compared to post-transcriptional purification methods. As such, our study aimed to reduce dsRNA formation during transcription by evolution of T7 RNAP. Through a combination of directed evolution and semi-rational design, we identified several beneficial mutations. These mutations were then randomly combined, resulting in a T7 RNAP variant that more effectively reduces the generation of dsRNA.

## Result

### Designing the FADS procedure

To establish a screening system for T7 RNAP, we optimized a molecular beacon-based method to detect the products of IVT. This real-time monitoring approach has shown a positive correlation between the fluorescence of the probe and RNA yield[34]. We hypothesized that positioning the molecular beacon at the 3’-end of long mRNA could potentially enable the distinction of RNA conformation and integrity. **Figure 1A (right panel)** illustrates our design, including various single-stranded DNA sequences representing target products (run-off RNA: BP-20), dsRNA with varying lengths (BP-0, BP-5, BP-10, BP-15: numbers indicate bases that can complementarily bind to the molecular beacon), and the incomplete mRNA.

**Figure 1.**
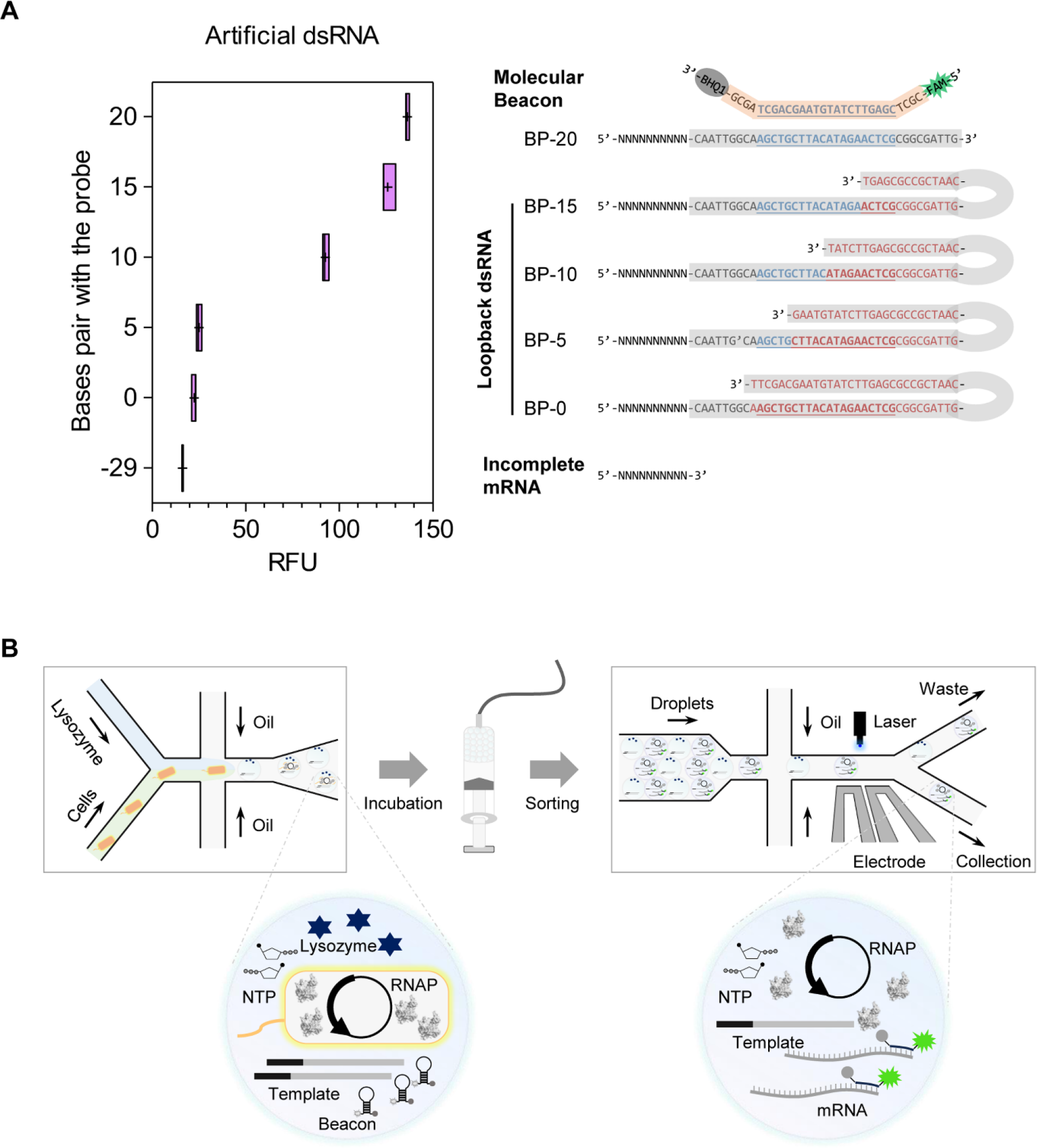
Overview of the FADS: **A. Relative fluorescence units (RFU) of the molecular beacon binding with ssDNA.** Equimolar single-stranded DNA was denatured, annealed, and incubated with molecular beacons at 37°C for 10 minutes. Then the fluorescence was measured (n=12). BP-20 represents the run-off RNA, which serves as the target product. BP-0, BP-5, BP-10, and BP-15 are designed to simulate dsRNA, where the numbers indicate the complementary bases on ssDNA that bind to the molecular beacon. “+” represents the mean value. **B. Generation and sorting of droplets.** The droplets contain all the necessary components for IVT, along with lysozyme and beacon. A dual aqueous phases chip channel was used to prevent premature cell lysis. Droplets were incubated in a syringe to facilitate lysozyme-mediated cell lysis and T7 RNAP-driven transcription after formation. Finally, the droplets were reinjected into a sorting chip for fluorescence detection and screening.

After incubating equimolar amounts of single-stranded DNA with the molecular beacon at 37℃, we observed a significant increase in fluorescence intensity in the reaction tubes as the number of complementary bases increased. Notably, the DNA lacking a complementary region, simulating incomplete mRNA, exhibited the lowest fluorescent after incubation **(Figure 1A, left panel)**. These results indicate that this method can effectively discriminate the conformation and integrity of single-stranded nucleic acids, making it suitable for mutant screening. In IVT reactions, theoretically, mutants with higher fluorescence intensity are expected to produce fewer dsRNA or/and incomplete mRNA.

For screening T7 RNAP, FADS was chosen to meet the high-throughput requirements of protein evolution. We validated the aforementioned probe-based method by utilizing purified T7 RNAP and *E. coli* strains BL21 expressing T7 RNAP. The fluorescence of reaction tubes showed a significant increase with higher enzyme or cell concentrations **(Figure S1A and Figure S1B)**. Moreover, when different lengths of DNA were used as templates, the signal-to-noise ratio of the system remained above 2 **(Figure S1C)**, indicating that the probe-based method is applicable for FADS. To prevent premature fragmentation of cells, we used a PDMS (polydimethylsiloxane) chip with dual aqueous phases, where lysozyme was mixed with the cells on the chip and exerted its activity within the droplets, as shown in **Figure 1B**. The number of droplets was controlled to be roughly ten times the number of cells, aiming to ensure that each microdroplet contained at most one cell. Droplets were collected into a syringe and incubated at 37℃ to facilitate cell lysis and RNA transcription. Subsequently, the droplets were introduced into a sorting chip, and only positive droplets exhibiting fluorescence above the threshold were collected for further investigation.

### FADS-based screening of T7 RNAP

We generated several random mutant libraries with a diversity ranging from 10^5^ to 10^6^ **(Table 1)**, using error-prone PCR. T7 RNAP expression was induced in each library, and droplets were generated for further analysis. These droplets were incubated to facilitate lysozyme-mediated cell lysis and T7 RNAP-driven transcription. To determine the sorting threshold, droplets encapsulating *E. coli* expressing wild-type T7 RNAP were prepared using the same procedure and incubation conditions.

**Table 1.** Parameters for the screening of L3 and L5 by FADS.

| Libraries | Lib3 |  |  | Lib5 |  |  |
| --- | --- | --- | --- | --- | --- | --- |
| Diversity | 5.7×10<sup>5</sup> |  |  | 2.6×10<sup>6</sup> |  |  |
| RFU | Round 1 | Round 2 | Mutant | Round 1 | Round 2 | Mutant |
| WT <sup>a</sup> | < 1.2 | < 2.5 | Mut1: V214A | < 2.7 | < 2.3 | Mut6: V186I |
| Threshold | 2.0 | 5.2 | Mut2: N233D | 4.1 | 5.7 | Mut7: F162S/A247T |
| Ratio | > 1.67 | > 2.08 | Mut3: A465T | > 1.52 | > 2.48 | Mut8: F416L |
| Count | 3539 | 122 | Mut4: A615T | 6793 | 204 | Mut9: L106P |
|  |  |  | Mut5: M369T |  |  | Mut10: D87G |

a. The maximum fluorescence of the wild-type serves as the threshold benchmark.

For the first round of mutant screening, the threshold was set at 1.5 times the maximum fluorescence of the wild-type, while for the second round, it ranged from 2 to 2.5 times the wild-type value. Considering the fluctuations in droplet fluorescence during each measurement, the screening threshold also varied and was typically determined by multiplying the value of the wild-type group obtained from the simultaneous test by a specific coefficient **(Table 1)**.

In the case of random library L3, 3539 and 122 positive droplets were collected after the first and second rounds of sorting, respectively, with fluorescence increases of 22.97% and 44.28% compared to the initial library **(Figure 2A)**. The gradual increase in fluorescence indicates an improved abundance of dominant variants in the library under equal cell count conditions. Similarly, library L5 yielded 6793 and 204 positive droplets, with fluorescence increases of 36.38% and 55.35%, respectively **(Figure 2B)**. The distribution of fluorescence is depicted in **Figure S2A and Figure S2B**. Following screening, a significant increase was observed in the proportion of droplets with high fluorescence, confirming the effectiveness of screening once again.

**Figure 2.**
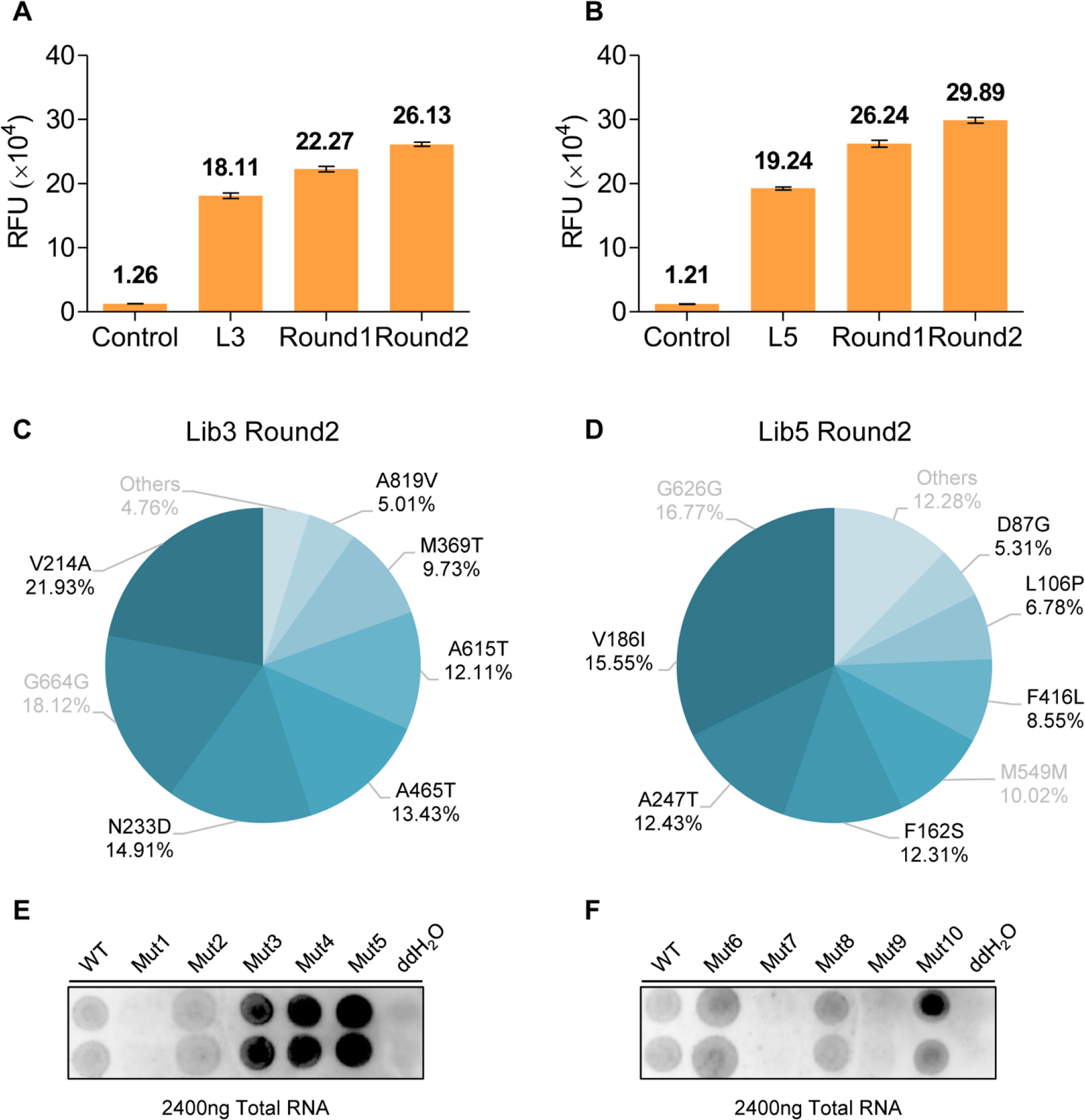
Random library screening and variants evaluation: **A (L3) and B (L5). RFU of IVT products from equivalent bacterial pellets.** DNA from positive droplets used for next-round library (Round1 and Round2). The *E. coli* libraries cultured for protein expression and 0.05OD cells were collected. After cell disruption, the fluorescence generated by the IVT was detected (n=3). **C (L3) and D (L5). Distribution and proportions of mutations.** Following two rounds of screening, DNA within the positive droplets was amplified via PCR and subsequently analyzed for sequencing using NGS. Silent mutations and mutations with frequencies below 5% are indicated in gray font. **E and F. dsRNA immunoblots of variants.** The IVT products of variants were diluted and spotted onto nylon membranes, and the J2 antibody was used to identify dsRNA. Mutant information: Mut1 (V214A), Mut2 (N233D), Mut3 (A465T), Mut4 (A615T), Mut5 (M369T), Mut6 (V186I), Mut7 (F162S/A247T), Mut8 (F416L), Mut9 (L106P), and Mut10 (D87G). 5μL ddH2O was used as the negative control.

After PCR amplification, we conducted next-generation sequencing (NGS) on the DNA extracted from positive droplets. The single-point mutations with a frequency exceeding 5% are illustrated in **Figure 2C and Figure 2D**. Among them, V214A (21.93%) and V186I (15.55%) were the most frequent amino acid mutation types in their respective libraries. We also identified several high-frequency silent mutations, such as G664G in L3 and G626G in L5, indicating that these sites may serve as hotspots during error-prone PCR. Interestingly, Sanger sequencing revealed a double-mutant variant with F162S and A247T, which explains why the frequencies of F162S and A247T mutations in L5 are closely matched.

We isolated clones and purified the proteins containing the mentioned sites, obtaining T7 RNAP variants named Mut1 to Mut10 **(Table 1)**. Subsequently, we analyzed the dsRNA content in their transcription products. As shown in **Figure 2E and Figure 2F**, the dsRNA content of Mut1, Mut7, and Mut9 was significantly lower than that of the wild-type, confirming the validity of our screening. However, Mut9 exhibited a transcriptional activity only 0.1% of wild-type, resulting in extremely low RNA production. Coincidentally, Mut2 showed an approximate 8% improvement in RNA integrity, while its activity was substantially lower than wild-type **(data not shown)**. These two variants may require additional evolution steps to enhance their activity and RNA yield.

It is perplexing that the products of Mut3, Mut4, and Mut5 exhibited significantly higher dsRNA content **(Figure 2E)**. We speculate that this may be due to false-positive results of FADS, or these variants exhibited increased production of another source of dsRNA during transcription, namely the dsRNA dependence on complementary pairing[35]. Although we screened T7 RNAP with reduced dsRNA production using FADS successfully, the enrichment of low- yield variants like Mut9 and Mut2 highlights the need for stricter monitoring of yield during the screening.

### Semi-rational design of T7 RNAP

Due to limited data, our structural and energetic analysis of Mut1 and Mut7 did not provide conclusive insights into the mechanism behind dsRNA reduction. Therefore, we decided to explore potential sites in spatially adjacent regions of F162, V214, or A247. On the other hand, we hypothesize that facilitating conformational transitions of T7 RNAP can significantly decrease the formation of abortive RNAs and ultimately reduce the dsRNA by-products[35]. An effective approach to achieve this goal is by reducing the free energy of the elongation conformation. This ensures a smoother transition of conformation and enhances the stability of the elongation complex, thus favoring the formation of run-off RNA. As described in the Materials and methods section, we conducted a semi-rational design exploration.

Through a comprehensive analysis of the protein structure and calculations using FoldX and PROSS **(Table S1)**, 43, 47, 70, 180, 189, 228, 237, 238, 243, 743, and 747 were selected as preliminary candidates **(Figure 3A and Figure 3B)**. Among these positions, 43, 47, 70, 180, and 189 are located in regions with significant conformational changes, while 228, 237, 238, 243, 743, and 747 are in the intercalating hairpin or specificity loop, adjacent to V214 and A247. Additionally, PROSS suggested the site 383 as a candidate **(Table S1)**, which is located in the non-template DNA strand-binding region. It believed to stabilize the non-template DNA and enhance the processivity of T7 RNAP, thus reducing incomplete RNA. Since mutations at 43 and 47 have already been proven to be beneficial for reducing dsRNA[36,37], we finally created ten single-site (A70, K180, D189, S228, V237, G238, T243, A383, I743, and L747) saturated libraries and employed the traditional microtiter plate method for screening. The selection of a saturated library aims to avoid potential functional impairments of site-directed mutagenesis caused by the limitations of simulation calculations.

**Figure 3.**
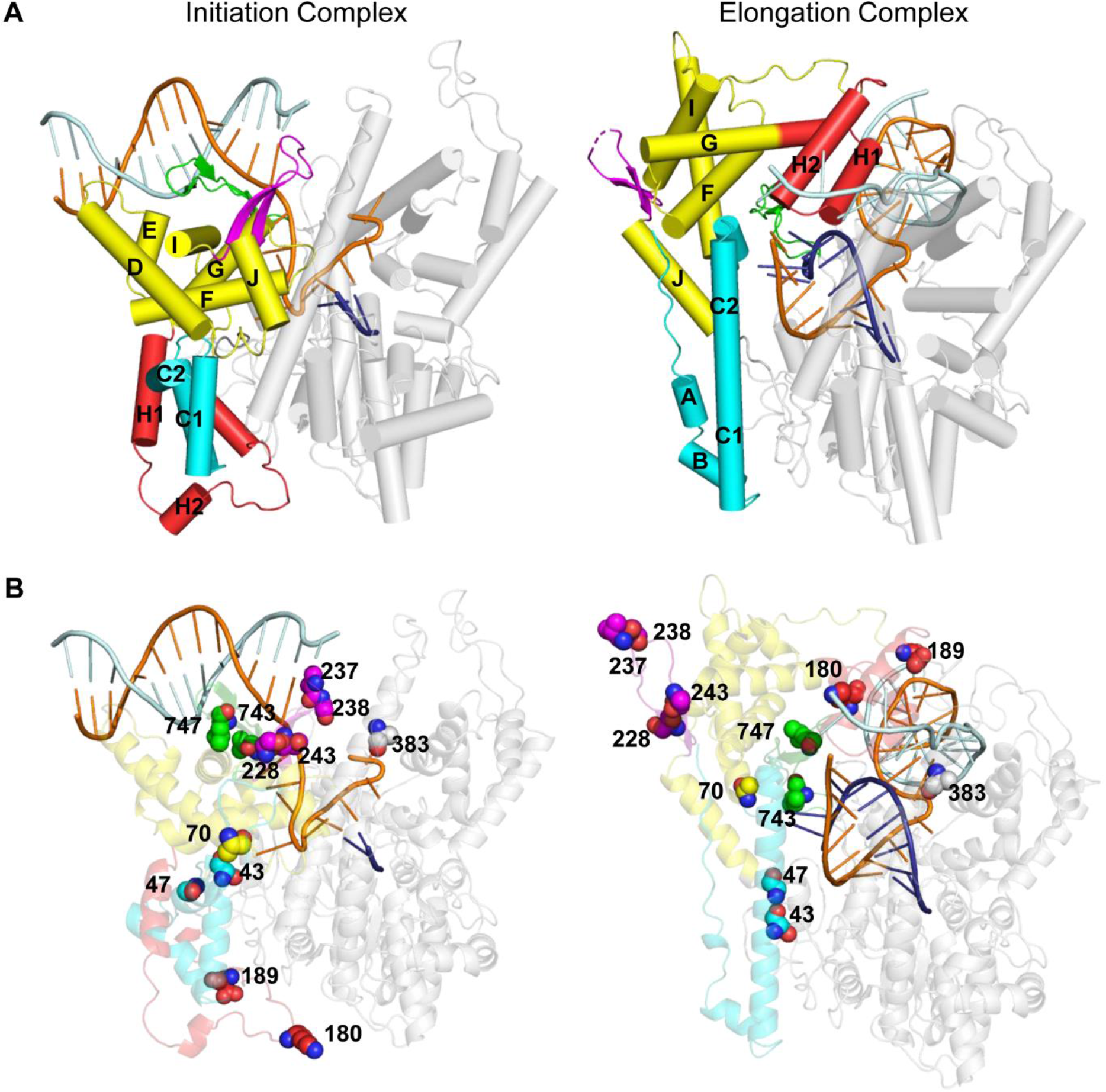
Structure-guided semi-rational design for decreased dsRNA: **A. Comparison of the structures of the T7 RNAP Initiation (left, PDB: 1QLN) and Elongation (right, PDB: 1MSW) complexes.** The molecules have been similarly oriented by superposition of their palm domains. The C-terminal domain is shown as gray transparent cartoon to allow views of the DNA and RNA. The N-terminal domain is colored as follows: promoter binding domain is yellow, the C-helix is cyan, the subdomain H domain is red, the specificity loop is green, the intercalating hairpin opens the double DNA is magenta. The template DNA is shown in orange, non-template DNA in pale cyan and RNA in deep blue. **B. Locations of the recommended positions.** Recommended positions are shown as spheres. The missing gaps in both the initiation and elongation complexes were repaired using Modeller. Proteins and substrates are colored as A. The protein is rendered as a transparent cartoon to highlight the recommended positions. The figure was generated using the PyMOL software.

To ensure a comprehensive evaluation of each variant, we improved the molecular beacon screening system. As shown in the upper right inset of **Figure 4A**, the 3’-end FAM probe is consistent with the function of beacon used in FADS, which is designed to screen mutants with reduced loopback dsRNA or incomplete RNA. The Cy5 probe positioned at the 5’-end of the RNA allows for the analysis of the yield of RNA[34], and perhaps provides an indication of the molar ratio between abortive RNA and run-off RNA. Sorting mutants based on the FAM to Cy5 ratio in each well ensures the enrichment of high-yield variants with significantly reduced dsRNA levels. We analyzed the fluorescence ratio changes over time for Mut1 and Mut10, observed that the ratios across various wells tended to converge and stabilize after 60 minutes. This indicates that the probes have annealed to the mRNA in a stable manner, and the fluorescence ratio at this point can represent the quality of the products. **(Figure S2 A and Figure S2 B)**.

**Figure 4.**
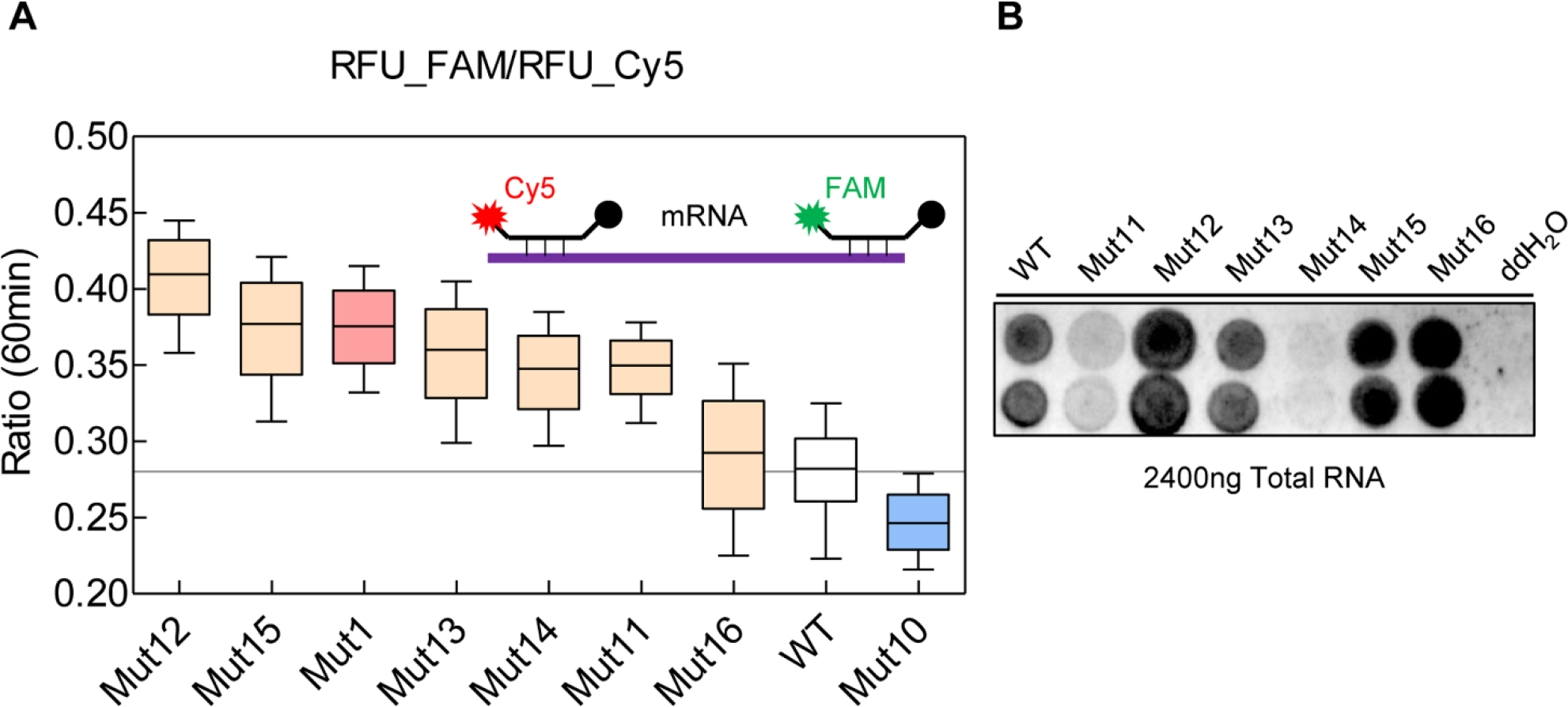
Microtiter plate screening and variant evaluation: **A. Dual probe detection of RNA products.** The purified T7 RNAP mutants were used for IVT, and the fluorescence of Cy5 and FAM was measured after 60 minutes. After a round of primary screening, each candidate was re-inoculated into a microplate for secondary screening (n>6). The superior mutants were identified by ranking them based on the FAM/Cy5 ratio. The fluorescence ratio of Mut1 and Mut10 serves as positive and negative controls, respectively. The black line marks the wild-type ratio of 0.2802, and any values above it is considered candidate mutants. The illustrations in the top-right corner indicate the probe positions and fluorescence types, with the 3’-end FAM probe designed to screen for mutants with reduced loopback dsRNA or incomplete RNA, and the Cy5 probe allowing for analysis of RNA yield and the molar ratio between abortive RNA and run-off RNA. **B. dsRNA immunoblots of variants.** The IVT products of variants were diluted and spotted onto nylon membranes, and the J2 antibody was used to identify dsRNA. Mutant information: Mut11 (K180E), Mut12 (S228F), Mut13 (G238L), Mut14 (A70Q), Mut15 (A383H), and Mut16 (I743L). 5μL ddH2O was used as the negative control.

After screening over 1000 mutants, we identified six mutants with fluorescence ratios higher than the wild-type **(Figure 4A)**: Mut11 (K180E), Mut12 (S228F), Mut13 (G238L), Mut14 (A70Q), Mut15 (A383H), and Mut16 (I743L).

We then quantified the transcriptional products of these mutants. The total RNA yield of all six mutants did not significantly differ from the wild-type, indicating comparable overall transcription efficiency. Expectedly, the dsRNA content of Mut11 and Mut14 was significantly lower compared to the wild-type **(Figure 4B)**.

Similar to the FADS experiment, the screening results from the saturated library revealed certain variants, particularly Mut12 and Mut15, where the dsRNA content did not align with the fluorescence ratio ranking. This discrepancy implies the presence of additional factors during the screening process that were not considered. To capture a wider range of beneficial variants, we will lower the threshold during subsequent screening.

### Generating Novel Mutants via DNA Shuffling

To further optimize the T7 RNAP, the four variants with low dsRNA content while meeting production requirements were selected. These mutants had specific mutations at five sites: V214A, F162S, A247T, K180E, and A70Q. We then constructed a DNA shuffling library consisting of approximately 500 clones, encompassing a theoretical total of 120 mutation types. Through screening with microtiter plates, we identified a variant that exhibited lower dsRNA production compared to its parental T7 RNAP, named Mut17 (A70Q/F162S/K180E).

We subsequently analyzed the transcriptional products of Mut17 and its parental variants (Mut14, Mut11, and Mut7). As shown in **Figure 5**, all four variants exhibited significant reductions in dsRNA production during transcription compared to the wild-type. Mut11 exhibited approximately 7.28% of the wild-type’s dsRNA content, while Mut14 and Mut7 had dsRNA levels of 3.49% and 3.31%, respectively. Remarkably, Mut17, the second-generation mutant, showed significantly lower dsRNA content of only 1.80% and as low as 0.007 ng/μg in the solvent system of screening.

**Figure 5.**
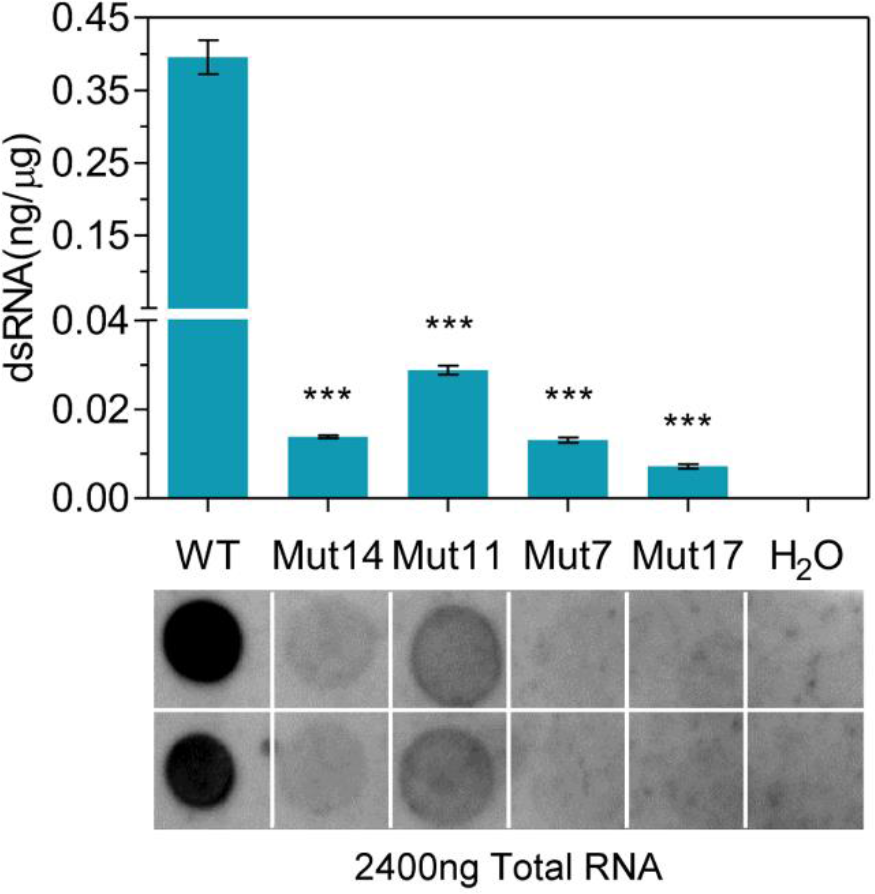
dsRNA content of Mut17 and its parental mutations: Upper: Quantification of dsRNA by ELISA, comparing wild-type and mutant variants (Mut14, Mut11, Mut7, and Mut17), based on standard dsRNA (composed of natural deoxyribonucleotides). N=3. *** indicates p < 0.0001. Lower: Immunoblots of dsRNA variants, with 5μL ddH2O as negative control. Mutant details: Mut14 (A70Q), Mut11 (K180E), Mut7 (F162S/A247T), Mut17 (A70Q/F162S/K180E).

### Immunogenicity of IVT products from mutants

Next, we evaluated the immunogenicity of the IVT products in murine RAW264.7 cells, and assessed the expression of EGFP mRNA in HEK293 cells. The 1kb dsDNA served as the template for T7 RNAP and its variants, containing the EGFP ORF. After introducing an equal amount of RNA, we detected the transcription and translation of IFN-β, a key effector in the immune response of cells against dsRNA[38]. As shown in **Figure 6A and Figure 6B**, IFN-β mRNA and protein were upregulated to varying degrees in RAW264.7 cells. The RNA synthesized by the wild-type T7 RNAP exhibited the most robust immune response, whereas the RNA derived from the mutants displayed a muted response. Specifically, the IFN-β mRNA of cells treated by Mut11 RNA was only 9.7% of the wild-type treated cells, and its protein was 12.93 pg/mL. With the exception of Mut11, which did not fully align with *in vitro* dsRNA quantification, the remaining mutants’ dsRNA content matched the cellular response, highlighting their effectiveness in reducing dsRNA **(Figure 5 and Figure 6)**. Moreover, the expression of EGFP showed no significant differences in quantity or quality among mRNAs generated by different T7 RNAPs **(Figure 6C)**, meeting the requirements for industrial applications.

**Figure 6.**
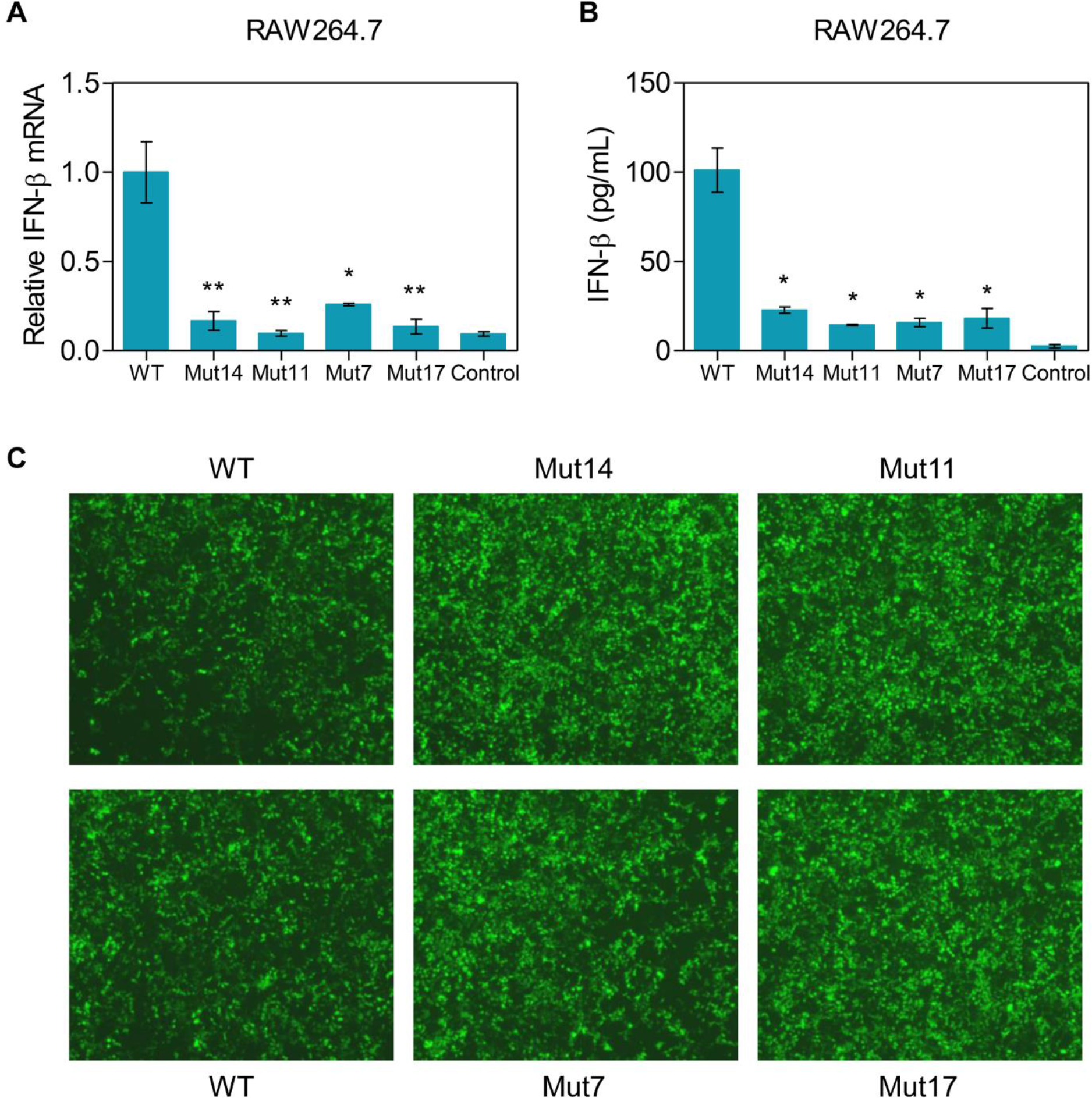
IFN-β response and EGFP expression in mammalian cells: **A. mRNA of IFN-β in murine RAW264.7 cells.** Cells were seeded in 24-well plates at a density of 0.5×10^5^ cells/well and transfected with 2 μg of each T7 RNAP transcript in triplicate. Following 24 h of incubation, total RNA was extracted, reverse transcribed, and analyzed for IFN-β mRNA, normalized to β-actin expression. Lipofectamine transfection served as a control. Significance levels: ** p < 0.005, * p < 0.05. **B. Quantification of IFN-β in murine RAW264.7 cells.** Cellular extracts were analyzed for IFN-β protein levels via ELISA using Mouse IFN-β as a standard. Lipofectamine transfection served as a control. Significance level: * p < 0.05. **C. Image of EGFP-expressing HEK293 cells.** HEK293 cells were transfected with EGFP mRNA using an identical protocol. After 24 h of incubation, cells were washed, diluted, and visualized under a fluorescence microscope.

Lastly, we conducted an evaluation of the transcriptional performance of Mut17, chosen as a representative variant, under varying conditions. **Figure S4** illustrates the tests performed on Mut17 with and without cap analogs, using diverse DNA templates. The results consistently demonstrated significantly lower dsRNA levels with Mut17, ranging from 0.18% to 1.62% of the wild-type. IVD-B is consistent with the solvent system of screening, exhibiting a dsRNA content of 1.62%, which closely matched the initial test result of 1.8% **(Figure 5)**. The incorporation of co-transcription capping and the use of modified nucleotides further reduced dsRNA production, with a minimum as low as 0.5pg/μg. However, the introduction of CleanCap® AU cap, especially when combined with N1-Me-Pseudo UTP, can affect the overall yield of RNA. Therefore, subsequent optimization might be necessary in such cases.

### RDRP and Terminal transferase activity of variants

To investigate the impact of mutations on reducing dsRNA, we conducted *in silico* studies on all these mutations. Five polymerases, including T7 RNAP wild-type and four variants, were simulated separately with ssRNA/ssRNA and dsDNA/mRNA **(Figure 7)**. The former simulated dsRNA formation, while the latter replicated T7 RNAP transcription. Since T7 RNAP does not require promoter recognition during RNA primer extension[21–25], we hypothesize that it adopts an elongation conformation when interacting with ssRNA/ssRNA substrates. Therefore, the dsDNA/mRNA substrate from 1MSW was modified to ssRNA/ssRNA as our structural model.

**Figure 7.**
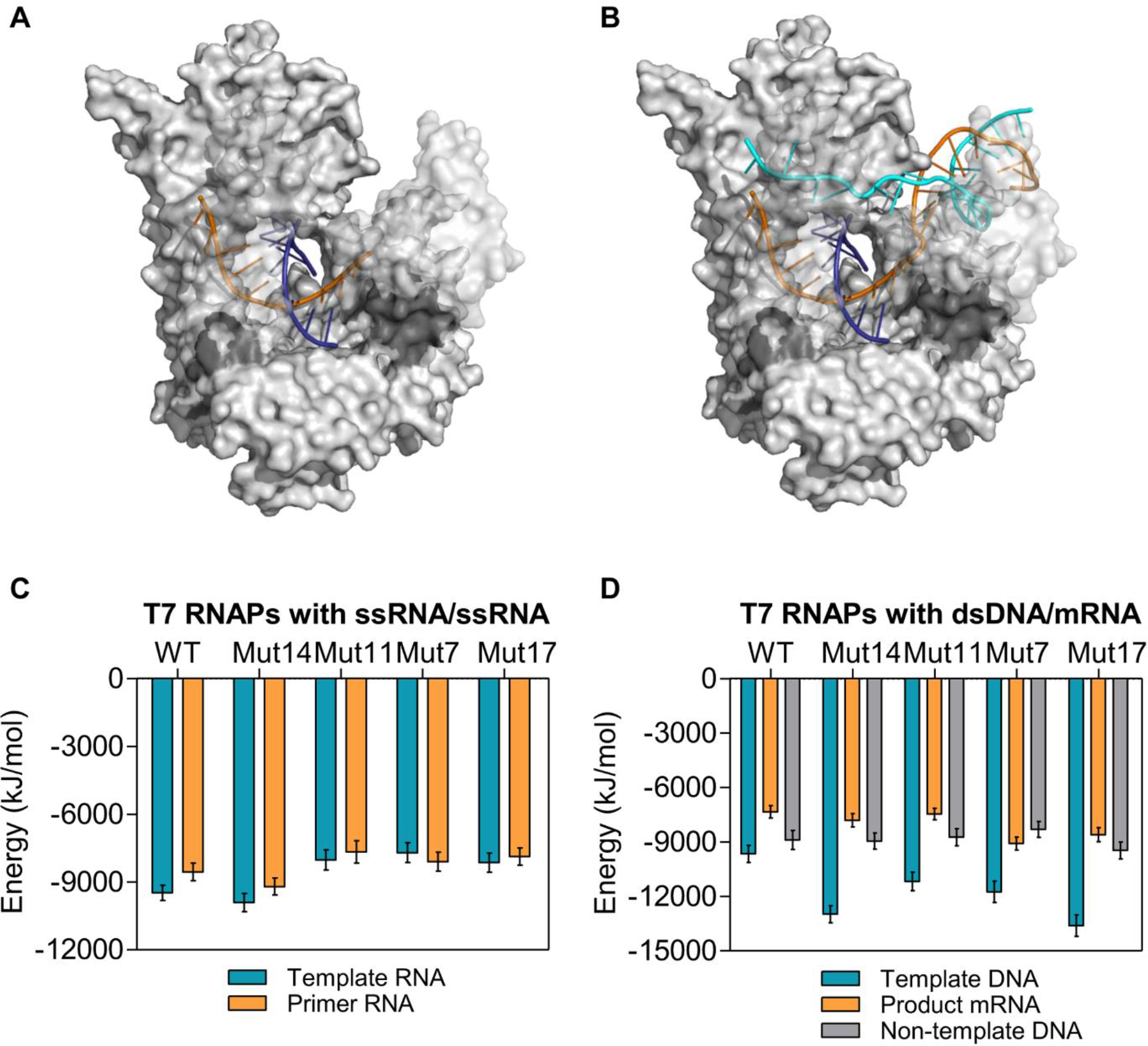
Molecular Dynamics Simulation of the interaction between T7 RNAPs and ssRNA/ssRNA (A and C) or dsDNA/mRNA (B and D): **A and B. Schematic diagram of the binding of T7 RNAP with ssRNA/ssRNA (A) or dsDNA/mRNA (B).** T7 RNAP is shown as a gray surface with residues 331-406 and 569-688 set to transparent to allow views of DNA and RNA. The template DNA or RNA are shown in orange, non-template DNA in pale cyan, mRNA or primer RNA in deep blue. The figure was generated using the PyMOL software. **C and D. The interaction energetics between T7 RNAPs and ssRNA/ssRNA (C) or dsDNA/mRNA (D).** All-atom MD simulations were performed using the GROMACS software package. The interaction energetics include electrostatic and Van der Waals contributions averaged from simulated trajectories.

We conducted a comparative analysis of protein-substrate interactions in all simulated systems. Our findings revealed that, except for Mut14 (A70Q), the variants exhibited higher binding energy with the RNA template when T7 RNAP bound to the ssRNA/ssRNA complex **(Figure 7A and Figure 7C)**. This suggests a clear indication of reduced RDRP activity in the variants, as they show a decreased tendency to utilize RNA as a template. Similarly, this may have had an impact on the terminal transferase activity of T7 RNAP[37]. In addition, we calculated the energy during the extension process of T7 RNAP. As expected, the mutants exhibited lower binding energy with DNA template compared to the wild-type **(Figure 7B and Figure 7D)**. However, there was no significant trend observed in the binding energetics with mRNA or the non-template DNA strand. Both ssRNA/ssRNA and dsRNA/mRNA systems clearly indicates the differences in template selection between the T7 RNAP wild-type and mutants.

To investigate these, we designed experiments to test the relative RDRP activity of T7 RNAP. The variants were diluted to the same transcriptional activity (i.e., DNA-dependent RNAP activity, DDRP), and then incubated with a hairpin structural chimeric substrate. As shown in **Figure 8**, under different concentrations, all 4 variants exhibited less than 50% of the fluorescence value compared to the wild-type. This signifies a lower efficiency in the formation of double- stranded products by the variants, implying a decrease in RDRP activity, likely responsible for the decrease in dsRNA content. However, it remains unclear why Mut14 shows a lower binding energy than the wild-type, while also reducing the synthesis level of dsRNA **(Figure 5 and Figure 7C)**.

**Figure 8.**
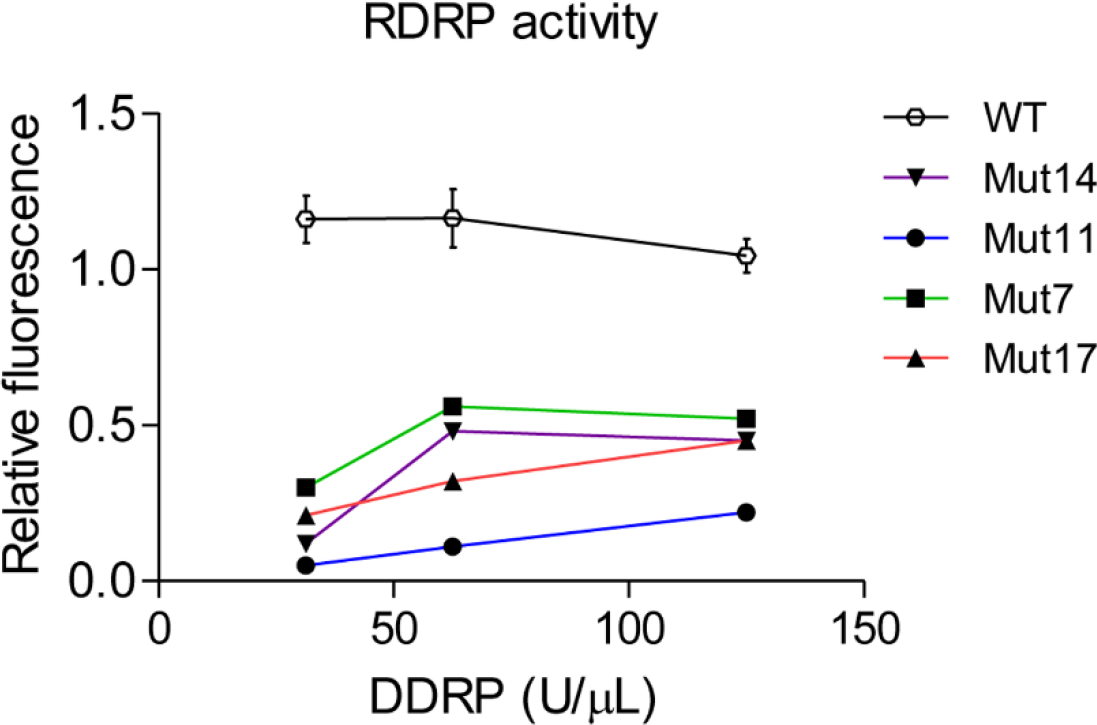
Relative RDRP activity: The hairpin oligonucleotide serves as a substrate to distinguish RDRP from DDRP activity of T7 RNAP. It forms a hairpin with a dC loop and RNA stem. Short poly(U) is complementarily paired with long poly(A) sequences to mimic a primer-template complex. T7 RNAP was diluted to 125 U/μL, 62.5 U/μL, and 31.25 U/μL for RDRP activity testing (n=3).

Subsequently, a 3’-end heterogeneity assay was employed to characterize the terminal transfer activity of T7 RNAP. We first transcribed RNA used mutants and wild-type T7 RNAP with DNA templates lacking a poly(A) sequence. Then, the *E. coli* poly(A) polymerase was utilized to add a poly(A) tail to the RNA. After reverse transcription, the nucleotide composition at the 3’-end was analyzed using NGS. **Figure 9** shows that all five T7 RNAPs produced a significant proportion of non-target RNA (n>0 or n<0). The most abundant among these were RNAs with an additional one or two nucleotides at the end (wild-type: n+2, others: n+1) **(Figure S5)**, confirming the strong terminal transferase activity of T7 RNAP.

**Figure 9.**
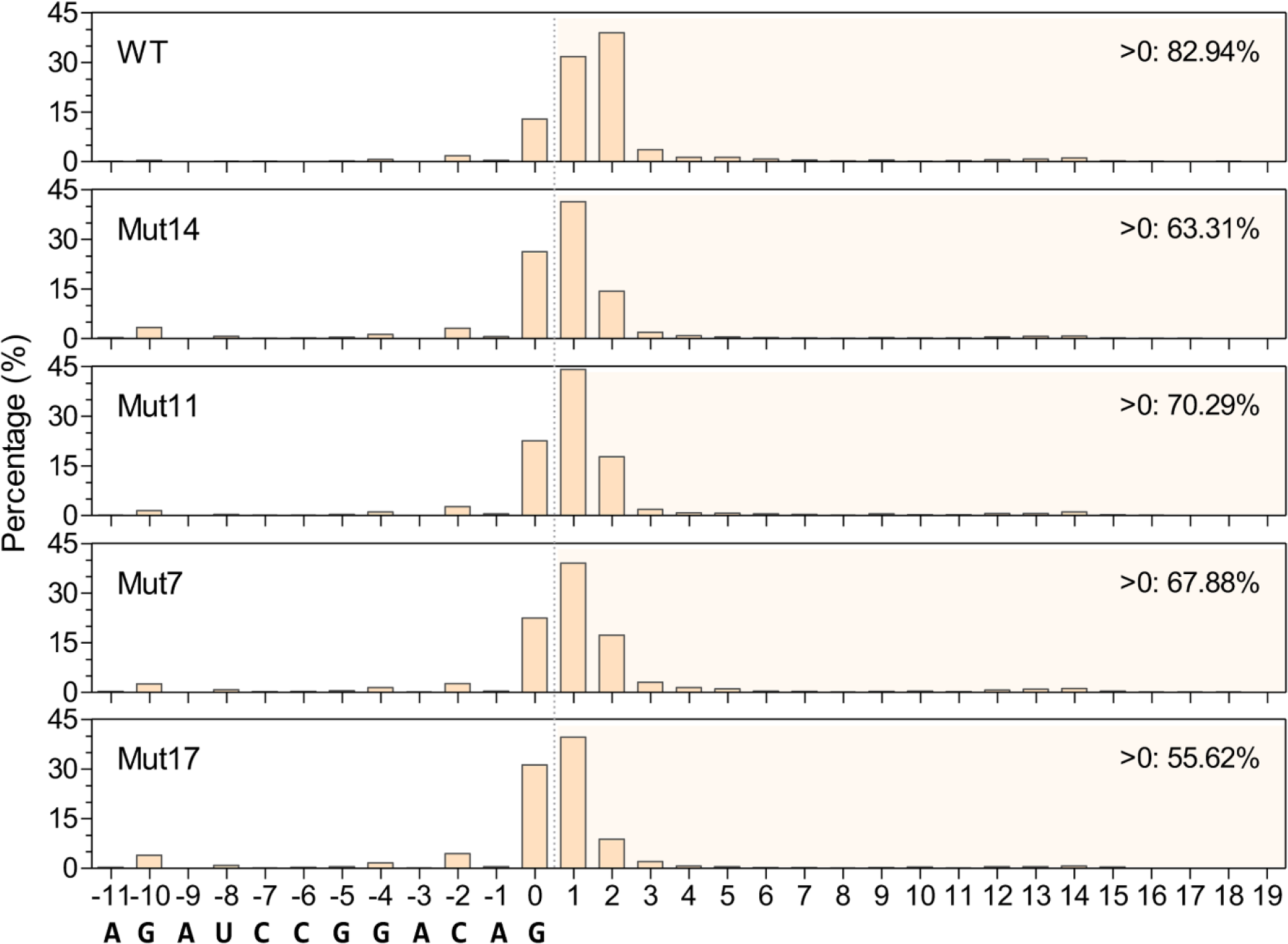
Heterogeneity in the 3’-end of RNA products: Reverse transcription was performed on poly(A)-tailed transcripts, followed by NGS analysis to assess the terminal base composition of each RNA. The bottom bases represent the 3’-end sequence of target RNA. Each graph includes numbers on the right side, indicating the percentage of RNA molecules that are longer than the DNA template. At the terminal of RNA, the addition of bases is random, and the numbers indicate the quantity of bases without specifying their types.

When focusing on the proportion of RNA with n>0, which reflects the terminal transferase activity, we found that these RNAs accounted for 82.94% in the wild-type, while the mutants exhibited the following percentages: Mut11 (70.29%), Mut7 (67.88%), Mut14 (63.31%), and Mut17 (55.62%) **(Figure 9)**. Importantly, these percentages are highly consistent with the abundance of dsRNA in the products (**Figure 5**). These results indicate a significant reduction in terminal transferase activity of mutants, which, together with the decrease in RDRP activity, contributes to the decrease in dsRNA. Specifically, the terminal transferase activity may play a more crucial role, as evidenced by the lowest accumulation of n>0 RNAs observed in Mut17, which also exhibits the lowest dsRNA production **(Figure 5 and Figure 9)**. We evaluated the major RNAs (**Figure S6**: n-1 to n+3) in the IVT products as an indicator of RNA integrity. The mutants’ major RNA showed a slightly lower percentage compared to the wild-type, attributed to an increase in n-10 and n-2 RNAs. However, as demonstrated in **Figure 6C**, minor losses of nucleotides likely have little effect on mRNA translation efficiency, potentially attributed to the UTR and poly(A) tail.

Considered that experimental validation establishes a significant correlation between RDRP and terminal transfer activities with the dsRNA content, screening strategies based on these activities are expected to provide new tools for further evolution of T7 RNAP.

## Discussion

In our study, we utilized a combination of FADS-based directed evolution and computer-aided semi-rational design, and obtained several T7 RNAP variants with significantly reduced dsRNA. Among these variants, Mut17 (A70Q/F162S/K180E) exhibited the lowest dsRNA content in the products. When co-transcription capping was performed using CleanCap® AU cap and N1-Me-Pseudo UTP, the dsRNA levels reached 0.5 pg/μg. The testing of RDRP and terminal transferase activity revealed the mechanism behind the decreased dsRNA in the variants’ products, as both activities closely associated with dsRNA formation demonstrated a significant decrease[35]. Specifically, the decrease in terminal transferase activity of variants appears to be the primary factor contributing to the reduction of dsRNA. This is supported by the strong positive correlation observed between this activity and dsRNA production among the different mutants **(Figure 5 and Figure 9)**. Based on the presented results, screening strategies focusing on the terminal transferase or RDRP activity would be a hopeful choice for further evolution of T7 RNAP.

Residue A70 belongs to the transition region between helix C and helix D, in close proximity to helix J and helix F. It is precisely positioned within the RNA exit channel constituted by Helix D, Helix J, and the specificity loop **(Figure 3A and 3B)**. The Mut14 mutation likely stabilizes the C/D helices, thus stabilizing the elongation conformation complex. Interestingly, Mut14 (A70Q) exhibits lower binding energy to RNA templates, potentially enhancing RDRP activity. However, in practice, both the RDRP and terminal transferase activities of Mut14 are reduced **(Figure 8 and Figure 9)**, which contributes to the decrease of dsRNA during IVT. This may imply the presence of more intricate factors influencing the RDRP activity of T7 RNAP. F162 and K180 reside on two helices of subdomain H, contributing to the stability of the transcription bubble during transcription **(Figure 3)**. While A247T is located in the transition region between the intercalating hairpin and J-helix. These three residues do not make direct contact with the template chain, yet simulations reveal stronger DNA template binding and weaker RNA template binding compared to the wild- type. This indicates that residues lacking direct interactions with the template can also alter the preference of T7 RNAP for the template. In an IVT system, there exists a competitive relationship between the transcriptional activity and the RDRP activity of the polymerase. When the polymerase prefers to bind to dsDNA/mRNA substrates, the activity of RDRP is weakened. Therefore, either enhancing the binding to DNA templates or reducing the binding to RNA templates is conducive to reducing the production of dsRNA during the *in vitro* transcription.

Before the design of the FADS procedure, we noted that even the wild-type T7 RNAP produced a minimal amount of dsRNA **(Figure 5)**, which significantly impedes FADS screening due to the difficulty in enriching candidates based on such minimal quantitative variations. However, in non-denaturing gel electrophoresis, we observed that dsRNA comprised about one-third to one-half of total RNA after IVT, identify by the acridine orange[39]. We concluded that FADS and this electrophoresis identify all RNA molecules containing complementary regions, which are present in a considerable quantity. The reduction of such RNAs caused by the decrease in dsRNA is sufficient to distinguish mutants from the wild-type. While J2 antibodies recognize and bind to the complementary regions in RNA, and the more complementary bases there are, the more antibodies bind.

On the other hand, the molecular beacon method was initially designed for real-time monitoring of T7 RNAP transcription, where the fluorescence reflects RNA yield[34]. However, during our FADS screening, low-yield variants, such as Mut2 and Mut9, were unexpectedly enriched. We hypothesize that when using long templates, specifically 4k or 9k in our study, the fluorescence generated by beacons targeting the 3’-end reflects only the amount of run-off RNA, not the total RNA yield. Consequently, variants with lower yields but improved integrity or reduced dsRNA levels, such as Mut2 and Mut9, have the potential to be identified **(Figure 2)**. Fortunately, the dual-probe screening has successfully addressed this limitation, ensuring an acceptable yield of the generated mutants **(Figure 4)**. This method will also be applied to FADS by expanding the fluorescence channels of the equipment.

In the random library L6, we identified a pseudo-positive variant, Mut18, which accounted for a high proportion of 86% after two rounds of screening **(data not shown)**. Mut18 contained five mutations at positions K8N/Q232L/N233D/E242D/N488D. We observed that variant Mut18 displayed significantly reduced transcriptional (DDRP) activity, which can be attributed to mutations at positions 232/233/242 that affect the functionality of the “intercalating hairpin” during promoter recognition or template unwinding[40]. However, when incubated alone with the probe, it markedly triggers the fluorescence of FAM. This indicates that the molecular beacon can elicit fluorescence not only through its binding to the target mRNA but also via direct interaction with Mut18. We suspect that Mut18 exhibits a higher affinity for hairpin structural probes compared to the T7 promoter, leading to their unwinding and subsequent fluorescence activation. This further prompted us to develop alternative screening strategies based on the direct activity of T7 RNAP.

## Materials and methods

### Oligonucleotides and reagents

Synthetic oligonucleotides were obtained from Sangon Biotechnology. NTP solutions, carbenicillin, pyrophosphatase, and Murine RNase inhibitor were acquired from Yeasen Biotechnology. Anhydrotetracycline, inorganic salt, and glycerol were purchase from Sigma-Aldrich.

### Beacon testing applied to artificial dsRNA

A total of 30 pM of artificial dsRNA **(Figure 1A)** was mixed with 500 nM molecular beacon and 2 μL of 10 × Transcription Buffer (CAT#: 10670ES, Yeasen Biotechnology). The volume was then adjusted to 20 μL with ddH2O. The mixture was incubated at 95℃ for 5 minutes and placed on ice for 10 minutes. Fluorescence was measured using a SpectraMax iD3 Microplate Detection System (Molecular Devices) with excitation/emission wavelengths of 480/525nm (Cutoff 520nm).

### FADS

To construct a random library, error-prone PCR was employed as previously described[41]. The Mn^2+^ concentration was 0.1 mM, and *E. coli* DH10B-Plus (CAT#: DE1072, Weidi Biotechnology) was chosen as the host. After overnight culture, the library was inoculated into fresh LB medium at a 2% ratio and incubated at 37℃ for 2 hours. To induce the expression of T7 RNAP, 200 ng/mL anhydrotetracycline was added to the culture, followed by further incubation at 16℃ for 16 hours. After harvesting, the pellet corresponding to 1OD was washed with 1 × Transcription Buffer and resuspended. For liquid phase A, a mixture was prepared by combining 5 μL of the bacterial suspension with 95 μL of 1 × Transcription Buffer and 20 mM NTPs. Liquid phase B was prepared by dissolving 1 μg DNA template, 1 μM beacon, and 0.2 mg/mL lysozyme in 100 μL of 1 × Transcription Buffer. Using the DREM cell droplet generation and sorting system (TmaxTree Biotechnology), liquid phases A and B were mixed in a 1:1 ratio on a PDMS chip, subsequently generating droplets with an approximate diameter of 25 μm. The same procedure was employed to prepare droplets containing *E. coli* expressing wild-type T7 RNAP. Both sets of droplets were incubated at 37℃ for 2.5 hours to ensure complete cell lysis and in-droplet transcription reactions. The droplets containing the library were sorted and positive droplets were collected. The sorting threshold was set as 1.5-2.5 times the maximum fluorescence intensity observed in the wild-type droplets **(Table 1)**. A double volume of emulsion breaker (Drop-breaker, FluidicLab) was added to facilitate phase separation. The upper aqueous phase was then extracted for PCR amplification, and the products were used for sequencing and construction of a new library.

### Purification of T7 RNAP and variants

The plasmids were separately transformed into *E. coli* BL21 (CAT#: EC1001, Weidi Biotechnology), and cells were collected after protein expression. T7 RNAP and the variants were purified using Ni-NTA chromatography in 20 mM Tris-HCl pH 7.5, 100 mM NaCl, 10% Glycerol, and 10 mM Imidazole. The purified samples were then dialyzed overnight against 100 mM Tris-HCl pH 7.9, 100 mM NaCl, 2 mM EDTA, 10 mM DTT, 0.1% Triton X-100, and 50% Glycerol. In the microplate screening, the same purification procedure is used, but with a reduced volume of around 4 mL.

### *In vitro* transcription assays

Gradient dilutions of variants, wild-type, and control (CAT#: 10626ES, Yeasen Biotechnology) T7 RNAPs were used to assess relative activity based on their *in vitro* transcription RNA yield. The IVT of T7 RNAP was performed following the procedure of the T7 High Yield RNA Synthesis Kit (CAT#: 10623ES, Yeasen Biotechnology). Briefly, a mixture of T7 RNAP, 1 μg DNA template, 10 mM NTPs, and 2 μL of 10× Transcription Buffer was prepared, and the volume was adjusted to 20 μL with DEPC-treated water. The reaction mixture was incubated at 37℃ for 2 hours, and the RNA was precipitated using LiCl. The pellet was washed twice with pre-chilled 70% ethanol and subsequently dissolved in DEPC-treated water. The resuspended RNA was quantified using a NanoDrop™ One (ThermoFisher Scientific). 250 U of T7 RNAP were added typically for IVT, aimed at comparing the performance of mutants. For co- transcription capping experiments, an additional 10 mM of CleanCap^®^ Reagent AU (CAT#: N-7114, TriLink Biotechnology) or CleanCap^®^ Reagent AG (3’OMe) (CAT#: N-7413, TriLink Biotechnology) analog was added.

### dsRNA immunoblots

A total of 2.4 μg of RNA from each variant were spotted onto Nytran SuPerCharge (SPC) blotting membranes (Cat#: 10416216, Cytiva) and dried at 80℃ for 40 minutes. The membranes were subsequently blocked in TBS-T buffer (20 mM Tris, pH 7.4, 150 mM NaCl, 0.1% (v/v) Tween-20) containing 5% (w/v) non-fat dried milk. For the detection of dsRNA, the membranes were incubated with Mouse monoclonal antibody J2 (Cat#: ab288755, Abcam) at a dilution of 1/1000 for 1 hour at room temperature. After washing 4-6 times, the membranes were immersed in TBS-T buffer containing Peroxidase-conjugated Donkey Anti-Mouse IgG (H+L) at a dilution of 1/2000 and incubated for 1 hour at room temperature. Following another round of washing (4-6 times), Super ECL Detection Reagent (CAT#: 36208ES, Yeasen Biotechnology) was used for visualization, and the images were recorded using the ChemiDoc MP Imaging System (Bio-Rad).

### Semi-rational design

The structure of initiation complex from a crystal structure (PDB: 1QLN), and elongation complex from a crystal structure (PDB: 1MSW). We removed waters and repaired the missing gaps (residues 1-5, 56-71 in the 1QLN and residues 1, 233-240, 364-374 in the 1MSW) used Modeller[42,43]. The free energy differences between the folded and unfolded structures of both the initiation and elongation conformations were predicted using a global PositionScan with FoldX[44], and these differences were compared to those of the wild-type (ΔΔG). Additionally, PROSS was used to predict mutations that are beneficial to the stability of the elongation conformation[45]. Structural analysis was performed using the visualization software PyMol[46].

### dsRNA ELISA

To quantify the dsRNA in the IVT products, a Double-stranded RNA (dsRNA) ELISA kit (CAT#: 36717ES, Yeasen Biotechnology) was used for testing the RNA samples. Briefly, standard substance and RNA samples were diluted to appropriate concentrations. 100 μL of each dilution was added to the reaction wells and incubated at room temperature for 1 hour. After discarding the supernatant, 100 μL of 1× Detection Antibody was added to each well and incubated at room temperature for another 1 hour. The samples were then discarded and the wells were washed. 100 μL of 1× Streptavidin-HRP was added to each well and incubated at room temperature for 30 minutes. TMB Substrate and Stop Solution were subsequently added, mixed, and the absorbance values were measured using the SpectraMax iD3 Microplate Detection System (Molecular Devices), under the condition of Ex = 450nm / Em = 650nm. A standard curve was generated to calculate the dsRNA content of each sample.

### Molecular dynamics simulation

The three-dimensional structure of the wild-type T7 RNAP was obtained by repairing the missing amino acids in the crystal structure (PDB: 1MSW) using the Modeller software. The mutations of T7 RNAP was obtained by mutating the wild-type structure using BIOVIA-Discovery Studio Visualizer (BIOVIA Corp). The dsDNA/mRNA substrate was derived from the ternary complex substrate of the crystal structure (PDB: 1MSW). The structure of the ssRNA/ssRNA was modified from the dsDNA/mRNA substrate with BIOVIA-Discovery Studio Visualizer: fist remove the non- template DNA, and then modified the DNA template to RNA strand, while retaining the 10 nucleotides at the 3’-end. The final ssRNA/ssRNA had a template strand of 5’-UUCGCCGUGU-3’ and a primer strand of 5’-GACACGGCGA- 3’. All molecular dynamics simulations were performed using the GROMACS-2022.1 software package[47–50], using the AMBER99SB force field with PARMBSC0 nucleic acid parameters and the Tip3p water model[51–53]. The steepest descent algorithm was used for energy minimization, and the Particle-Mesh Ewald method was used to treat long- range electrostatic interactions[54]. The ion concentration to neutralize the system were maintained 0.15M NaCl. The cutoff distance of van der Waals force (vdW) and short-range electrostatic interaction was set to 10 Å. The system temperature was maintained at 310K, with the neighbor list updated every 10 steps, a time step length of 2fs, and a MD simulation of 50ns.The interactions between protein and substrate were calculated from simulated trajectories using the gmx energy module in Gromacs. The interactions between protein and substrate energy includes Van der Waals forces and electrostatic interactions. The ele and vdW interaction energies were calculated at a cut-off distance of 40 Å.

### RDRP activity

T7 RNAP and variants were diluted to the following concentrations: 125 U/μL, 62.5 U/μL, 31.25 U/μL, respectively, based on their activities (DDRP activity). For each dilution, 1 μL of the enzyme was added to the previously mentioned TVT system. A single-stranded chimeric nucleotide chain (5’- AAAAAAAAAAAAAAAAAAAAdCdCdCdCdCUUUUUUUUUU-3’) at a concentration of 1.25 μM was used as the transcription template, which forms a stable hairpin structure after incubation at 65℃ for 2 minutes followed by 4℃ for 2 minutes. During a 10-minute incubation at 37℃, the T7 RNAP extended the 3’-end of the chimeric nucleotide chain, resulting in the formation of a complete double-stranded product. The reaction was then quenched by incubating at 65℃ for 5 minutes. According to the protocol of Picogreen dsDNA Quantitation Reagent (Cat#: 12641ES, Yeasen Biotechnology), a double-stranded dye was added to each reaction well. The fluorescence was measured using the SpectraMax iD3 Microplate Detection System (Molecular Devices) with excitation at 450nm and emission at 650nm to determine the relative RDRP activity for each variant.

### Heterogeneity analysis of the RNA 3’-end

To add a poly(A) tail to the 3’-end of RNA, *E. coli* poly(A) Polymerase (CAT#: 14801ES, Yeasen Biotechnology) and 1mM ATP was added to the samples and incubated at 37℃ for 30 minutes. Tailed products were then subjected to reverse transcription using M-MLV (H-) Reverse Transcriptase (Cat#: 11300ES, Yeasen Biotechnology) with a poly(T) primer, followed by PCR amplification of cDNA for sequencing.

### Transfection of cells

HEK293 and murine RAW264.7 cells were plated in Culture Flask (Cat#: 84022ES, Yeasen Biotechnology) and incubated for 36 h. After treatment with trypsin-EDTA, the cells plated in 24-well plates at a density of 0.5 × 10^5^ cells per well. Following the Lipofectamine 3000 Reagent (ThermoFisher Scientific) protocol, cells were grown to approximately 90% confluence before transfection. For transfection, 2 μg of capped RNA transcribed from different variants was dissolved in 50 μL of Opti-MEM Medium along with 4 μL of P3000 reagent. The solution was thoroughly mixed with diluted Lipofectamine 3000 Reagent at a 1:1 ratio and incubated at room temperature for 15 minutes. Subsequently, 50 μL volumes of different lipofectamine/RNA solutions were added to the wells of the three kinds of cells. The cells were then incubated at 37°C for 24 hours. After incubation, the cells were washed with PBS and released by treatment with 0.5% Trypsin-EDTA. The HEK293 cells were used for EGFP fluorescence analysis, while murine RAW264.7 cells were utilized for quantification of IFN-β mRNA and protein.

### Reporter cell line immunogenicity assays

The immunogenicity of RNA from each variant was tested in murine RAW264.7 cells, as previously described[36]. The qPCR primers for mouse β-actin and IFN-β are as follows: Mactin-F: 5’-AACAGTCCGCCTAGAAGCAC-3’, Mactin-R: 5’-CGTTGACATCCGTAAAGACC-3’; MIFN-F: 5’-GCCTTTGCCATCCAAGAGATGC-3’, MIFN-R: 5’-ACACTGTCTGCTGGTGGAGTTC-3’. The quantification of IFN-β protein was performed using the IFN-β/IFNB ELISA Kit (Cat#: CSB-E04945m, Huamei Biotechnology), following the manufacturer’s protocol.

