## Supplementary Information for "FADS and semi-rational design modified T7 RNA polymerase reduced dsRNA production, with lower terminal transferase and RDRP activities"

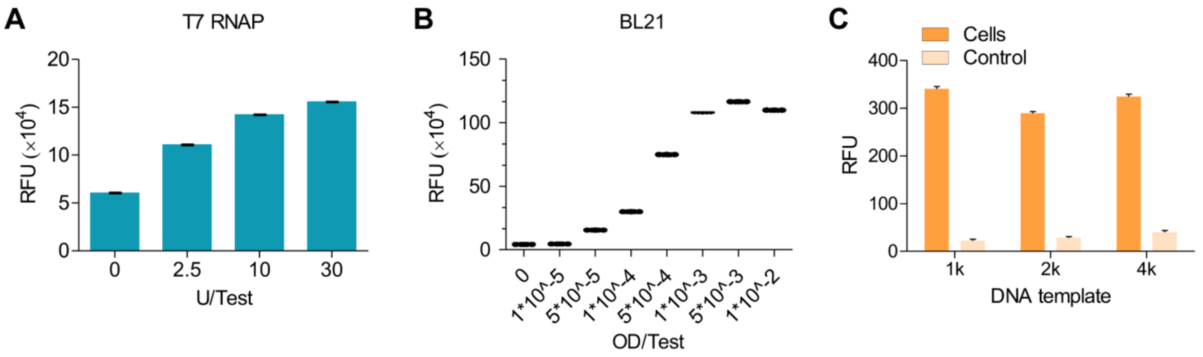


**Figure S1. Relative fluorescence units of the molecular beacon under different conditions.**

**A.** IVT using purified T7 RNAP.

**B.** IVT using BL21 expressing T7 RNAP.

**C.** IVT using purified T7 RNAP with DNA templates of various lengths.

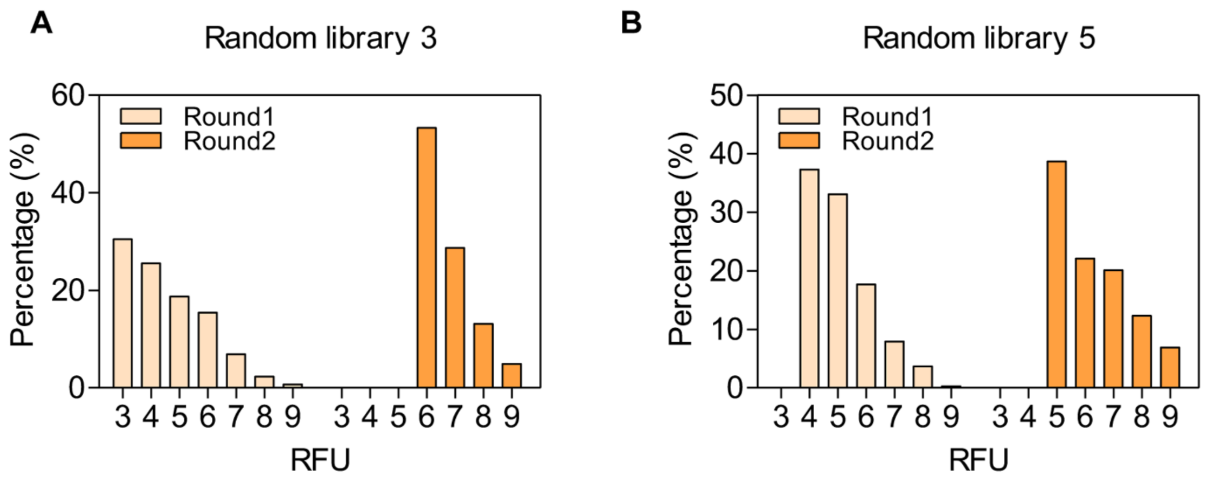


**Figure S2. The distribution of droplets with different RFU.**

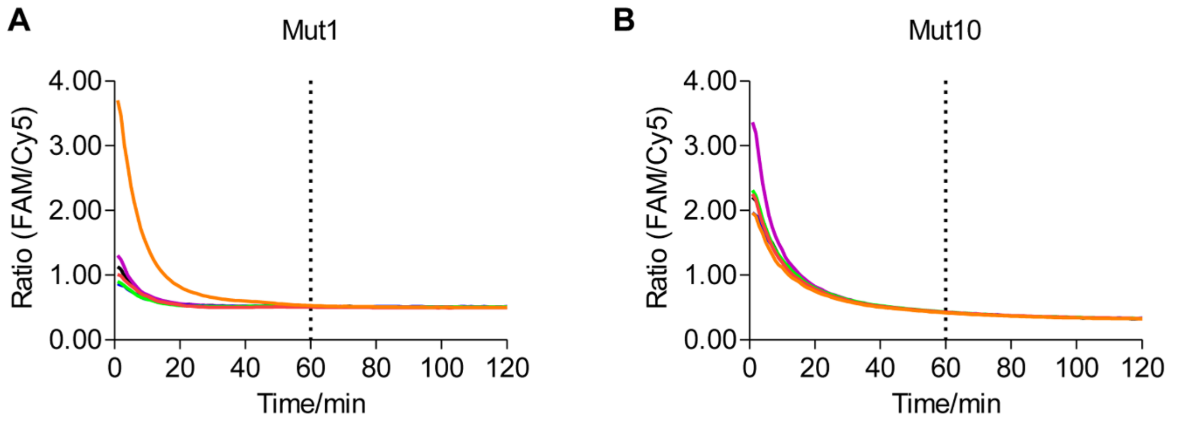


**Figure S3. The temporal trend of fluorescence ratios for Mut1 (A) and Mut10 (B).**

The fluorescence was collected every 30 seconds, and the data were obtained from six replicate wells for each mutant.

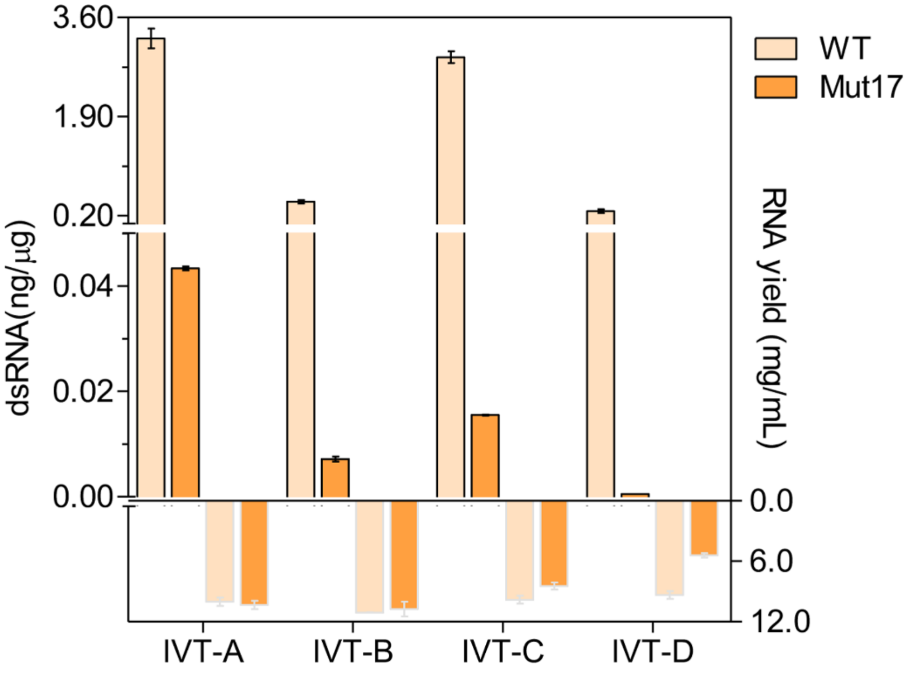


**Figure S4. The applicability of Mut17 and wild-type T7 RNAP.**

The left axis represents the level of dsRNA, while the right axis represents the RNA yield. IVT-A: DNA template with the natural T7 promoter (5'-TAATACGACTCACTATAGG-3') in the absence of cap analogs. IVT-B: DNA template with the optimized promoter (5'-TAATACGACTCACTATAAG-3') in the presence of m7(3'OMeG)(5')ppp(5')(2'OMeA)pG cap. IVT-C: DNA template with the optimized promoter (5'-TAATACGACTCACTATAGAT-3') in the presence of m7G(5')ppp(5')(2'OMeA)pU cap. IVT-D: Transcription using N1-Me-Pseudo UTP based on condition IVT-C.

**Most abundant**

Encoded: GUGCGGAAGAUCCGGACAG

| WT | | | |
|---|---|---|---|
| | 33.8% | GUGCGGAAGATCCGGACAGCC | n+2 |
| | 26.0% | GUGCGGAAGATCCGGACAGC | n+1 |
| | 10.7% | GUGCGGAAGATCCGGACAG | n |
| | 3.0% | GUGCGGAAGATCCGGACAGG | |
| | 1.7% | GUGCGGAAGATCCGGACAGCT | |
| | 1.3% | GUGCGGAAGATCCGGACAGCCG | |
| | 1.2% | GUGCGGAAGATCCGGACAGT | |
| | 1.1% | GUGCGGAAGATCCGGAC | |
| | 1.1% | GUGCGGAAGATCCGGACAGCCC | |
| | 1.0% | GUGCGGAAGATCCGGACAGCTTCCGCACATCTC | |

| Mut11 | | | |
|---|---|---|---|
| | 36.8% | GUGCGGAAGATCCGGACAGC | n+1 |
| | 20.0% | GUGCGGAAGATCCGGACAG | n |
| | 14.5% | GUGCGGAAGATCCGGACAGCC | n+2 |
| | 3.0% | GUGCGGAAGATCCGGACAGG | |
| | 2.3% | GUGCGGAAGATCCGGACAGT | |
| | 1.7% | GUGCGGAAGATCCGGAC | |
| | 1.3% | GUGCGGAAG | |
| | 1.1% | GUGCGGAAGATCCGGACAGCT | |
| | 1.0% | GUGCGGAAGATCCGGACAGCG | |
| | 0.9% | GUGCGGAAGATCCGGACAGCTTCCGCACATCTC | |

| Mut14 | | | |
|---|---|---|---|
| | 33.2% | GUGCGGAAGATCCGGACAGC | n+1 |
| | 23.6% | GUGCGGAAGATCCGGACAG | n |
| | 10.9% | GUGCGGAAGATCCGGACAGCC | n+2 |
| | 3.2% | GUGCGGAAGATCCGGACAGT | |
| | 3.0% | GUGCGGAAG | |
| | 3.0% | GUGCGGAAGATCCGGACAGG | |
| | 2.3% | GUGCGGAAGATCCGGAC | |
| | 1.2% | GUGCGGAAGATCCGGACAGCG | |
| | 0.9% | GUGCGGAAGATCCGGACAGCT | |
| | 0.9% | GUGCGGAAGATCCGG | |

**Most abundant**

Encoded: GUGCGGAAGAUCCGGACAG

| Mut17 | | | |
|---|---|---|---|
| | 29.5% | GUGCGGAAGATCCGGACAGC | n+1 |
| | 28.8% | GUGCGGAAGATCCGGACAG | n |
| | 6.4% | GUGCGGAAGATCCGGACAGCC | n+2 |
| | 5.1% | GUGCGGAAGATCCGGACAGT | |
| | 3.6% | GUGCGGAAG | |
| | 3.4% | GUGCGGAAGATCCGGAC | |
| | 3.1% | GUGCGGAAGATCCGGACAGG | |
| | 1.2% | GUGCGGAAGATCCGG | |
| | 0.8% | GUGCGGAAGATCCGGACAGCG | |
| | 0.7% | GUGCGGAAGAT | |

| Mut7 | | | |
|---|---|---|---|
| | 31.4% | GUGCGGAAGATCCGGACAGC | n+1 |
| | 19.9% | GUGCGGAAGATCCGGACAG | n |
| | 13.9% | GUGCGGAAGATCCGGACAGCC | n+2 |
| | 3.4% | GUGCGGAAGATCCGGACAGG | |
| | 2.5% | GUGCGGAAGATCCGGACAGT | |
| | 2.3% | GUGCGGAAG | |
| | 1.8% | GUGCGGAAGATCCGGAC | |
| | 1.1% | GUGCGGAAGATCCGG | |
| | 1.1% | GUGCGGAAGATCCGGACAGCG | |
| | 1.0% | GUGCGGAAGATCCGGACAGCCG | |

**Figure S5. Comparison of the most abundant 3'-end heterogeneities.**

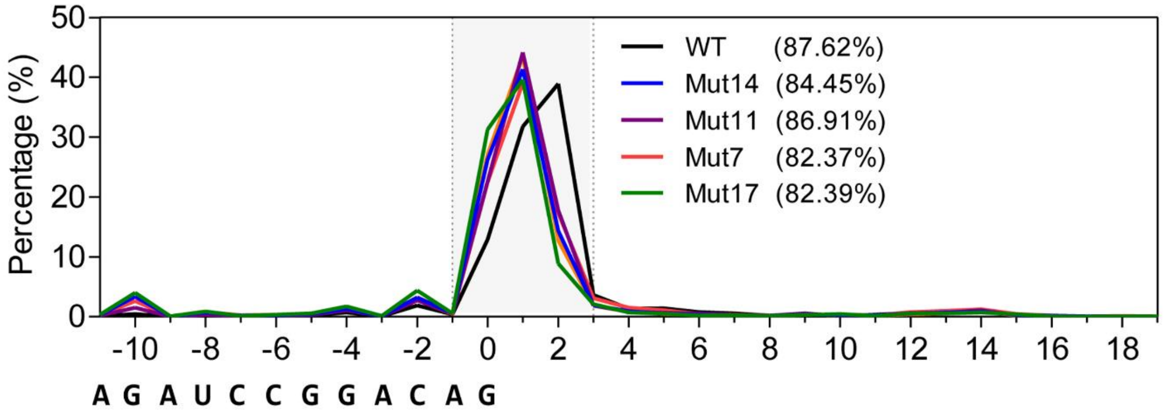


**Figure S6. The proportion of major products in the transcripts of each mutant.**
